# Tree-based quantification infers proteoform regulation in bottom-up proteomics data

**DOI:** 10.1101/2025.03.06.641844

**Authors:** Constantin Ammar, Marvin Thielert, Caroline A. M. Weiss, Edwin H. Rodriguez, Maximilian T. Strauss, Florian A. Rosenberger, Wen-Feng Zeng, Matthias Mann

## Abstract

Quantitative readout is essential in proteomics, yet current bioinformatics methods lack a framework to handle the inherent multi-level nature of the data (fragments, MS1 isotopes, charge states, modifications, peptides and genes). We present AlphaQuant, which introduces *tree-based quantification*. This approach organizes quantitative data into a hierarchical tree across levels. It allows differential analyses at fragment and MS1 level, recovering up to 50-fold more regulated proteins compared to a state-of-the-art approach. Using gradient boosting on tree features, we address the largely unsolved challenge of scoring quantification accuracy, as opposed to precision. Our method clusters peptides with similar quantitative behavior, providing a new approach to the protein grouping problem and enabling identification of regulated proteoforms directly from bottom-up data. Combined with deep learning classification, we infer phosphopeptides from proteome data alone, validating our findings with EGFR stimulation data. We then describe proteoform diversity across mouse tissues, revealing distinct patterns of post translational modifications and alternative splicing.

## Introduction

Mass spectrometry-based proteomics must both identify and quantify proteins across experimental conditions, time points, and disease states. While recent bioinformatic advances have substantially improved protein and peptide identification, quantification strategies still have considerable room for improvement. Modern data-independent acquisition (DIA) as well as established data-dependent acquisition (DDA) methods produce rich, structured datasets that enable direct comparisons via *label free quantification* (LFQ), making them particularly amenable to advanced computational analysis.

Bioinformatics approaches for LFQ fall into two main categories. The first encompasses reductionist approaches^1–7^ such as MaxLFQ^1^, which derive protein quantities by consolidating peptide, MS1, or fragment data while addressing challenges like varying ionization efficiency and missing values. Modern iterations like QuantUMS^6^ use machine learning to optimize quantification accuracy. While these approaches provide straightforward protein quantities for downstream analysis, they inevitably sacrifice some of the data’s complexity and richness. In response, a second category of methods focusses on differential quantification, preserving granular data in statistical analyses. These methods leverage peptide-level measurements^8–10^, peptide data enhanced with metadata^11,12^, and fragment-level information^13,14^, consistently demonstrating superior performance over traditional summarization-based methods by retaining detailed data structure. Additionally, they can perform statistics on missing values instead of relying on imputation^15,16^.

The quantification challenge is further complicated by *proteoform*^17^ diversity, where single genes generate multiple protein variants through alternative splicing or post-translational modifications (PTMs), opening up a largely unexplored dimension of proteomic complexity^18^. Top-down proteomics^19^, which analyzes intact proteins, offers the most direct approach to proteoform resolution, but faces technical limitations in protein separation and large ion analysis. Bottom-up proteomics, which analyzes proteolytically digested peptides, enables high-throughput, standardized measurements but introduces different challenges. Proteolytic digestion disconnects peptides of the same protein from each other, creating the protein grouping problem and complicating proteoform identification. Additionally, computational constraints typically restrict whole proteome analysis to unmodified peptides, relegating post-translational modification (PTM) analysis to specialized workflows. As many of these issues cannot be solved at the identification level, bioinformatics methods have proposed to use quantification for both protein grouping^20–22^ and proteoform detection^23–31^, exemplified by COPF^26^, FlexiQuant^27,28^ and PeCorA^30^. However, these tools are constrained to proteins with abundant peptide evidence^28^, optimized for diverse multi-condition datasets^26^, single peptide outliers^26,30^, and have not been integrated into standard differential analysis workflows.

We present AlphaQuant, an innovative end-to-end software pipeline that seamlessly integrates quantitative proteomics data with experimental design and acquisition parameters. At its core, AlphaQuant implements differential quantification down to the fragment level, preserving crucial information while enabling accurate modeling of quantitative noise for statistical comparisons between ions. To organize this complex data, we developed a novel *tree-based quantification* methodology that builds a hierarchical structure from fragments and MS1 isotopes up through charge states, modifications, peptides, and genes. This framework combines clustering algorithms with machine learning on tree-derived features to enable robust quantification and reveal proteoform groups through peptides showing coherent quantitative patterns. The versatility of AlphaQuant’s architecture makes it compatible with diverse quantification methods, with built-in support for the common search engines in both DDA and DIA workflows.

Here, we describe the conceptual underpinnings of tree-based quantification. We demonstrate how our method enhances differential quantification and provides a rigorous statistical solution for handling missing values. Using gradient boosting regression on tree-derived features, we establish a novel accuracy score that shows excellent classification performance. We validate AlphaQuant’s proteoform grouping capabilities using a custom experimental benchmark and apply it to two biological cases: detection of regulated phosphopeptides in EGFR-stimulated samples from unenriched proteomics data, and analysis of proteoform differences across eight tissues in mouse.

We provide AlphaQuant as an open source Python package, based on the scientific Python stack and the alphaX^32^ ecosystem, accessible through a Python API, Jupyter notebooks, or an easily installable graphical user interface.

## Results

### Tree-based quantification

Our algorithm begins with differential statistical analysis, which requires three key inputs: a search engine results table, a sample-to-condition mapping (e.g., samples 1-3 for case group, 4-6 for control), and comparison instructions. The method then conducts a differential comparison between these defined conditions. In case of multiple conditions, AlphaQuant can construct a median reference condition, against which all other conditions can be compared.

The tree structure of AlphaQuant is built around the *base ion* level, which is the most fundamental level of quantification. In DIA data, base ions can comprise both MS2 fragments and MS1 isotopes. Additionally, other quantitative measurements, such as precursor intensities, can serve as base ions. These base ions exhibit widely different intensity levels due to ionization efficiencies, isotopic distributions, and fragmentation patterns (**Fig. 1a**). Converting these intensities to log2 fold changes (log2FCs) between conditions enables direct comparisons, where base ions from the same peptide should show similar ratios between case and control, reflecting their common abundance change (**Fig. 1b**).

**Figure 1:**
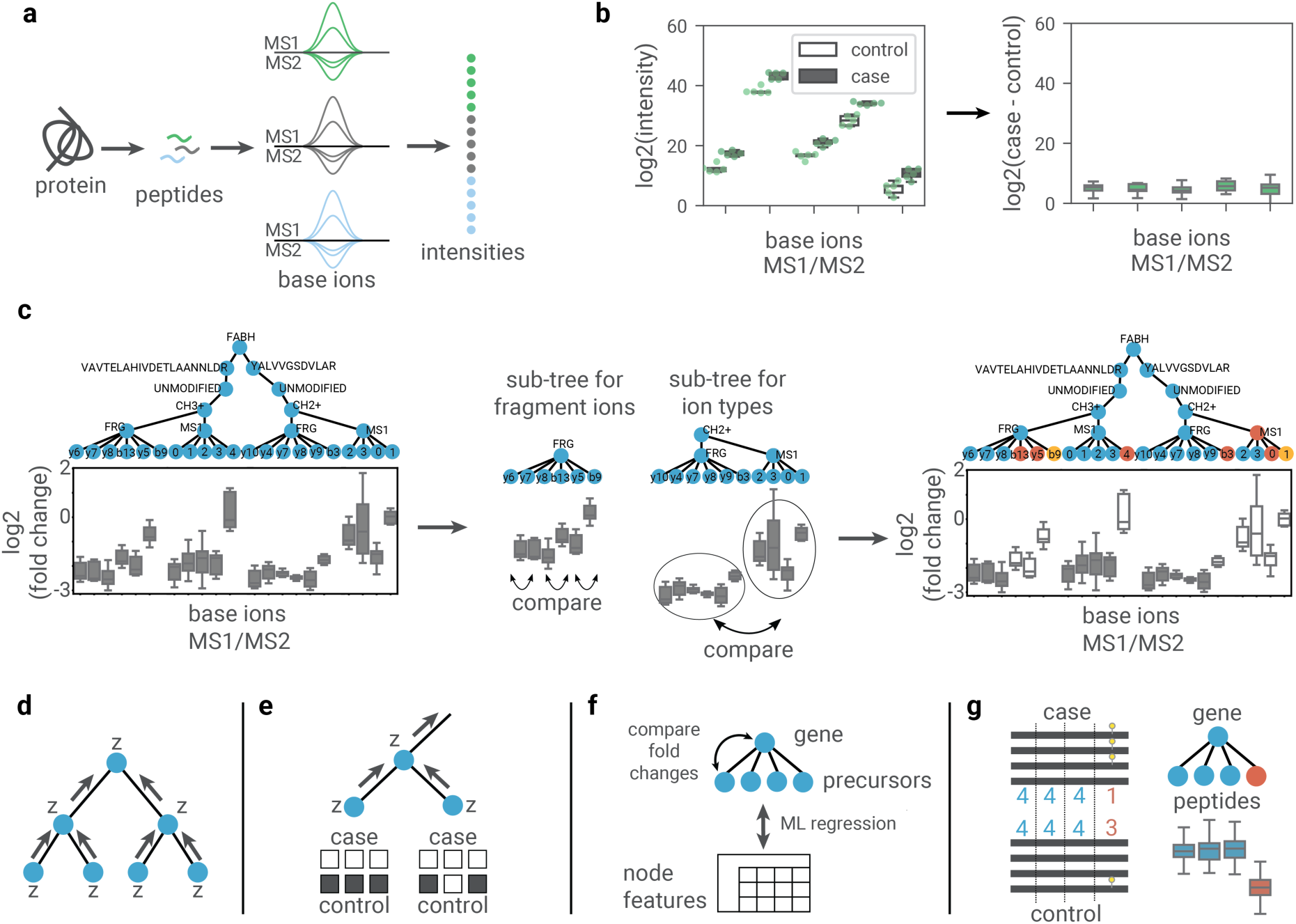
Main concepts of the tree-based quantification implementation termed AlphaQuant. a) Each protein quantified in a bottom-up proteomics experiment generates a multitude of quantitative features, such as fragments and MS1 isotopes ("base ions"). b) Base ions have different intensity levels and cannot be directly compared. Converting to log2 fold-changes between biological conditions creates directly comparable distributions. These distributions can be compared via a tailored statistical model. c) The fold-change transformed base ions are integrated into a hierarchical tree structure. Successive consistency checks proceed along the tree. Using hierarchical clustering, we assess the consistency among fragments, among MS1 isotopes, between fragments vs. MS1 isotopes, different charge states, modifications, and peptides, ultimately generating a clustered tree containing consistent (dark grey) elements. d) Propagation of Z-value transformed differential quantification results from base ion level along the tree. e) Statistical evaluation of precursors completely missing from one condition. f) Quality scoring of precursor nodes using gradient boosting regression. g) Detection of potential proteoforms by clustering peptides with similar quantitative behavior.

From the base ion level, we construct a hierarchical tree with nodes representing progressively higher organizational levels: ion type (fragments or MS1), charge state, modification status, peptide sequence, and ultimately the gene name (**Fig. 1c, left**). The lower tree levels from base ions up to the peptide sequence should show perfect consistency, as they all come from the same peptide. By using the gene name as node identifier, peptides from the same gene (including different isoforms) can be analyzed together and grouped by the clustering process.

To validate these expected consistencies, we perform statistical comparisons within each subtree level: between MS1 isotopes, charge states and fragments (**Fig. 1c, middle**). We model the null distribution of base ion log2FCs using empirical and replicate-based statistics^29^. This generates p-values to each base ion pair, allowing us to cluster ions together ions for which we cannot reject the null hypothesis of similar log2FCs. This method successfully identifies and removes inconsistent measurements, such as divergent fragment- and MS1 ions, retaining consistent base ions (**Fig. 1c, right**).

We perform differential quantification tests for all base ions of the tree using empirical and replicate-based noise distributions as described previously^9^. After transforming the results of each node into Z-values and propagating them along the tree, we correct for dependencies between base ions via appropriate covariance matrices (**Fig. 1d**).

We address missing values through counting statistics rather than imputation. When a protein is absent in one condition but quantified in another, we apply intensity-dependent binomial counting statistics per precursor to test the null hypothesis that the value is missing at random (**Fig. 1e**). Rejection of this test indicates that the protein is downregulated. These counting statistics are then integrated into the differential quantification as they propagate up the tree.

We enhance the clustered tree with additional properties such as peak shape quality, retention time, and identification q-values, reasoning that this information can be leveraged to evaluate quantification reliability. To condense this information into a single metric, we employ gradient boosting regression on features derived from precursor (defined as the modified, charged peptide sequence) nodes. For proteins with three or more precursors the regression model predicts the absolute deviation between protein and precursor log2FCs (**Fig. 1f**). This leverages the principle that protein values typically provide more accurate estimates than individual precursor values.

Feature combinations indicating high quality (e.g. consistent MS1 and fragments, high intensity, and good peak shape profiles) should correlate with smaller deviations.

Our approach enables proteoform inference by analyzing co-clustering peptide groups, since modifications and splicing create distinctive quantitative patterns for affected peptides. For example, consider a protein with four copies in two conditions, but varying phosphorylation: one copy phosphorylated in control and three copies phosphorylated in case (**Fig. 1g**). Here, the modified peptide shows a ratio of 3:1, while unmodified peptides maintain a 1:1 ratio. Tree-based quantification detects these distinctive patterns, with genes serving as tree roots to cover all isoforms.

### Differential quantification

To evaluate differential quantification of AlphaQuant, we utilized mixed-species datasets where organisms are combined at varying ratios across two conditions. As proteins are regulated according to their species of origin, this provided a ground truth for evaluating method performance, enabling straightforward calculation of true and false positive rates for each method (**Fig. 2a**).

**Figure 2:**
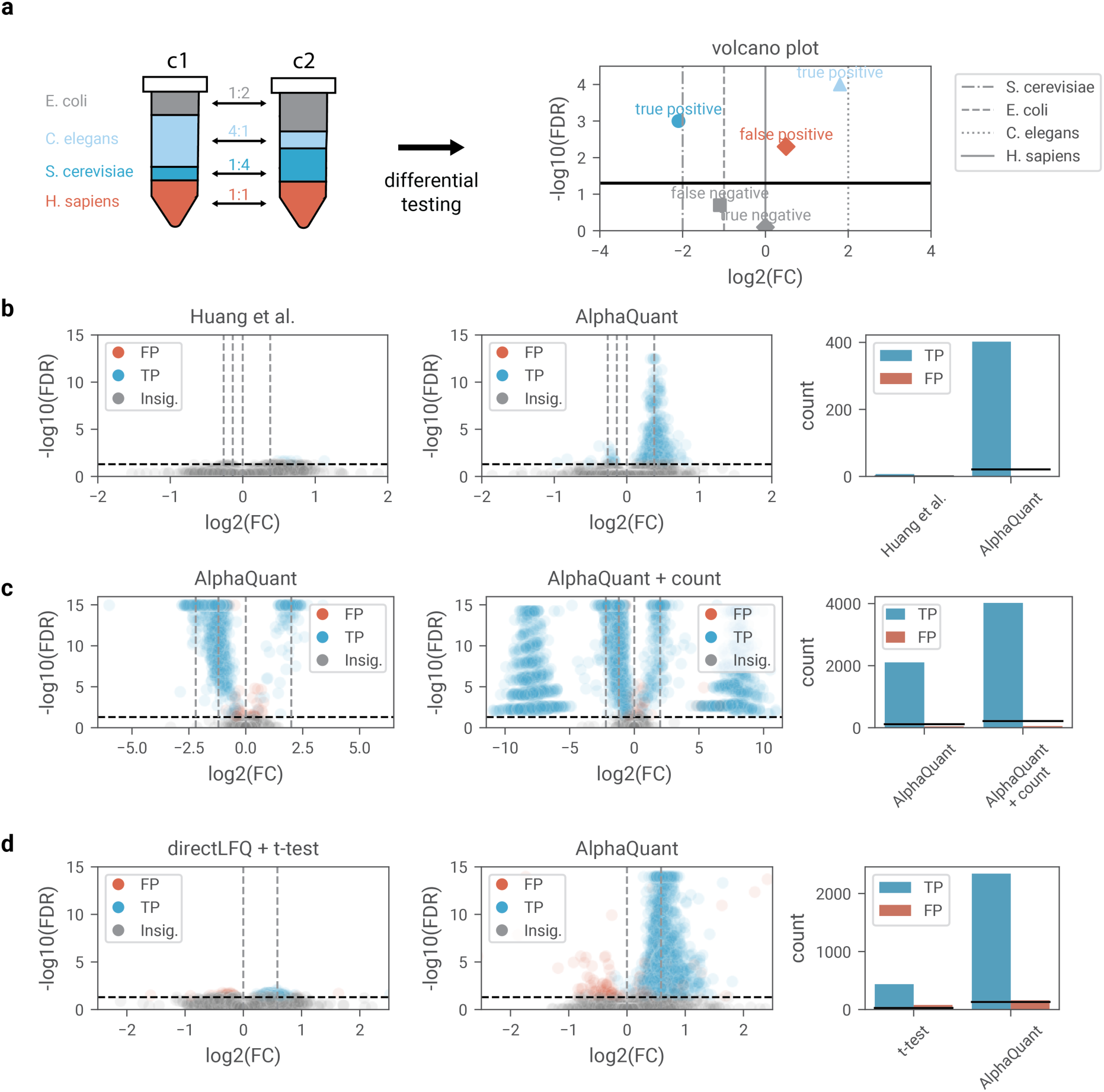
Benchmarking differential quantification on diverse benchmarking datasets. Black lines in the bar plots in panels b-d indicate the theoretically expected number of False Positives (FPs). a) Design of a mixed species experiment and classification of results into true positive (TP) and false positive (FP). b) Comparison a state-of-the-art method to AlphaQuant on a low log2FC benchmarking dataset. c) Impact of AlphaQuant’s counting statistics module for values that are completely missing in one condition. The two clouds of proteins with strong log2FCs on the right plot represent previously undetectable proteins recovered by the module. d) Comparison of a standard t-test analysis with directLFQ to AlphaQuant on a phospho-enriched benchmarking dataset.

We benchmarked AlphaQuant against a recent method^14^ that similarly combines MS1 and fragment-level quantification. In the most challenging dataset with subtle inter-species ratios of 30% or less, AlphaQuant identified several hundred regulated proteins—an over 50-fold improvement in detection sensitivity compared to this state-of-the-art approach (**Fig. 2b**). This dramatic increase in sensitivity stems from AlphaQuant’s tree-based model, which reaches high statistical significance when multiple fragments show consistent regulation patterns, thereby enabling reliable detection of subtle log2FCs. Importantly, this enhanced sensitivity does not compromise specificity, as AlphaQuant maintains well-controlled false discovery rates.

To further validate AlphaQuant, we analyzed a published high-ratio mixed-species dataset^14^, to investigate if tree-based quantification can improve detection of proteins with large log2FCs. While proteins with sufficient datapoints and strong FCs are typically quantified accurately by any method, those falling below detection limits in any condition create missing values. We first used AlphaQuant to quantify proteins that have a minimum of two replicate measurements in both conditions, which resulted in the successful recovery of nearly all regulated proteins as expected. We then expanded our analysis to proteins entirely absent from one condition using AlphaQuant’s intensity-based counting statistics module. This broader approach doubled the number of correctly identified regulated proteins, recovering almost all that were missing not at random with very good false discovery control (**Fig. 2c**). These results demonstrate that our combination of tree-based quantification and counting statistics provides a powerful method for detecting regulation of proteins despite missing values.

Given AlphaQuant’s enhanced sensitivity, we next evaluated its performance in differential quantification analyses of studies using PTM-enrichment. This is an important and particularly challenging case, which typically results in just one peptide that needs to be correctly quantified. We analyzed a previously published dataset, which had been designed to benchmark detection of phosphorylation in DIA data^33^. This dataset contains enriched phosphopeptides spiked into a HeLa background lysate at different ratios. Using AlphaQuant’s integrated site mapping and quantification workflow, we identified more than 2,500 regulated phospho-sites. In contrast, analysis of the mapped phospho-sites with our recently published directLFQ method^5^ combined with a t-test identified fewer than 500 sites (**Fig. 2d**). The false positives fall within the expected FDR range, but their systematic pattern of deviation suggests that the complex setup of spiking in phosphopeptides in the study may have introduced some systematic bias.

### Scoring quantitative accuracy

In quantitative proteomics, precision measures the reproducibility of measurements as quantified by the *coefficient of variation* (CV). While accuracy—the proximity of measurements to true quantities—is equally important, there are no straightforward metrics to assess it^34^.

We hypothesized that that we could infer accuracy using machine learning by leveraging the rich data available from tree-based quantification. Using a regression model trained on the absolute difference between the FC of the gene node and the FC of the precursor node, we developed a proxy for accuracy. This approach relies on two key premises: (i) when measuring many proteins, they will distribute around the true value, and (ii) a protein’s quantification, (approximated by the median value across all its precursors), provides a more accurate measurement than individual precursor measurements.

To benchmark accuracy, we generated a mixed-species dataset by spiking yeast proteins into a human background with a log2FC of 3.3. This ten-fold difference provided an unambiguous ground truth for accuracy assessment. We focused on the yeast subset and defined precursors as "outliers" if their measured log2FC deviates more than ±1 unit from the ground truth (i.e., measurements <2.3 or >4.3). Conversely, we classified precursors as "accurate" if their measured value within ±0.3 units of the true value (i.e., 3.0<x<3.6).

Applying AlphaQuant to the complete dataset, processed separately through both DIA-NN and Spectronaut, the AlphaQuant quality score (AQscore) readily distinguished between outliers and accurate precursors (**Fig. 3a,b**). The Area Under the Curve (AUC) values were 0.92 and 0.91 for DIA-NN and Spectronaut, respectively. This performance significantly exceeds that of individual quality scores provided by the search engines—an intended outcome since these scores serve as inputs to the AlphaQuant machine learning model, where they are integrated and analyzed alongside other performance metrics.

**Figure 3:**
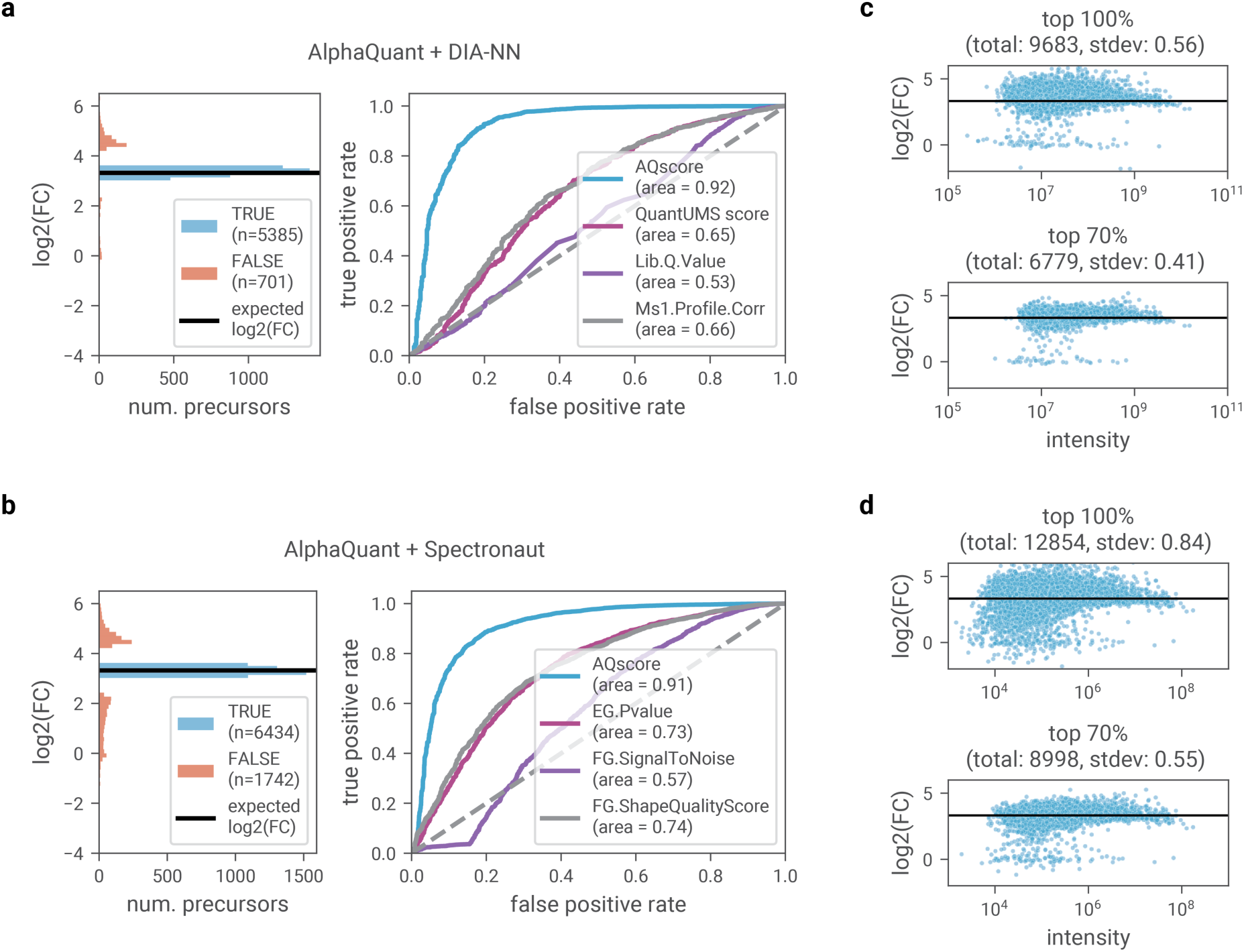
Evaluating quantitative accuracy at the precursor level. a) Left: Histogram showing the distribution of accurate precursors versus outliers. Right: ROC curves comparing the AlphaQuant accuracy score with quality scores provided by DIA-NN. b) Same analysis using Spectronaut processing. c) Distribution of yeast precursor processed by DIA-NN. d) Distribution of yeast precursors processed by Spectronaut.

For more granular analysis, we progressively filtered the data by removing the 30% of precursors with the lowest AQscores. The resulting ratio-intensity plots demonstrate how this filtering approach reduces the spread of precursors around the target value: from SD 0.56 to SD 0.41 in DIA-NN and from SD 0.84 to SD 0.55 in Spectronaut data (**Fig. 3c,d**). Points with clear alignment around zero likely are false identifications (human peptides identified as yeast peptides).

### Detecting regulated phosphopeptides from unenriched proteomics data

We hypothesized that outlier peptide detection could enable the inference of PTMs from unmodified peptides, as these show distinct quantitative behavior when a modified counterpart is present. We tested this concept with protein phosphorylation, a widespread PTM crucial in cellular signaling cascades.

We acquired a standard (non-phospho-enriched) proteomics dataset from HeLa cells, in which we compared control samples with those stimulated by Epidermal Growth Factor (EGF), known to trigger widespread phosphorylation responses^35^. Since the experimental design specifically targeted phosphorylation-driven changes, we considered phosphorylation to be the primary cause of outlier peptides. Examination of EGF receptor (EGFR) by AlphaQuant revealed five peptides that clustered separately. Sequence analysis using the AlphaMap^36^ package, which we integrated into AlphaQuant, revealed that they corresponded to previously documented and annotated phosphorylation sites (**Fig. 4a**). Notably, a significant subset of outlier peptides clustered within the autocatalytic C-terminal tail domain of EGFR, including the well-characterized autophosphorylation sites Y1172 and Y1197, demonstrating AlphaQuant’s ability to detect biologically relevant phosphorylation events without phospho-enrichment.

**Figure 4:**
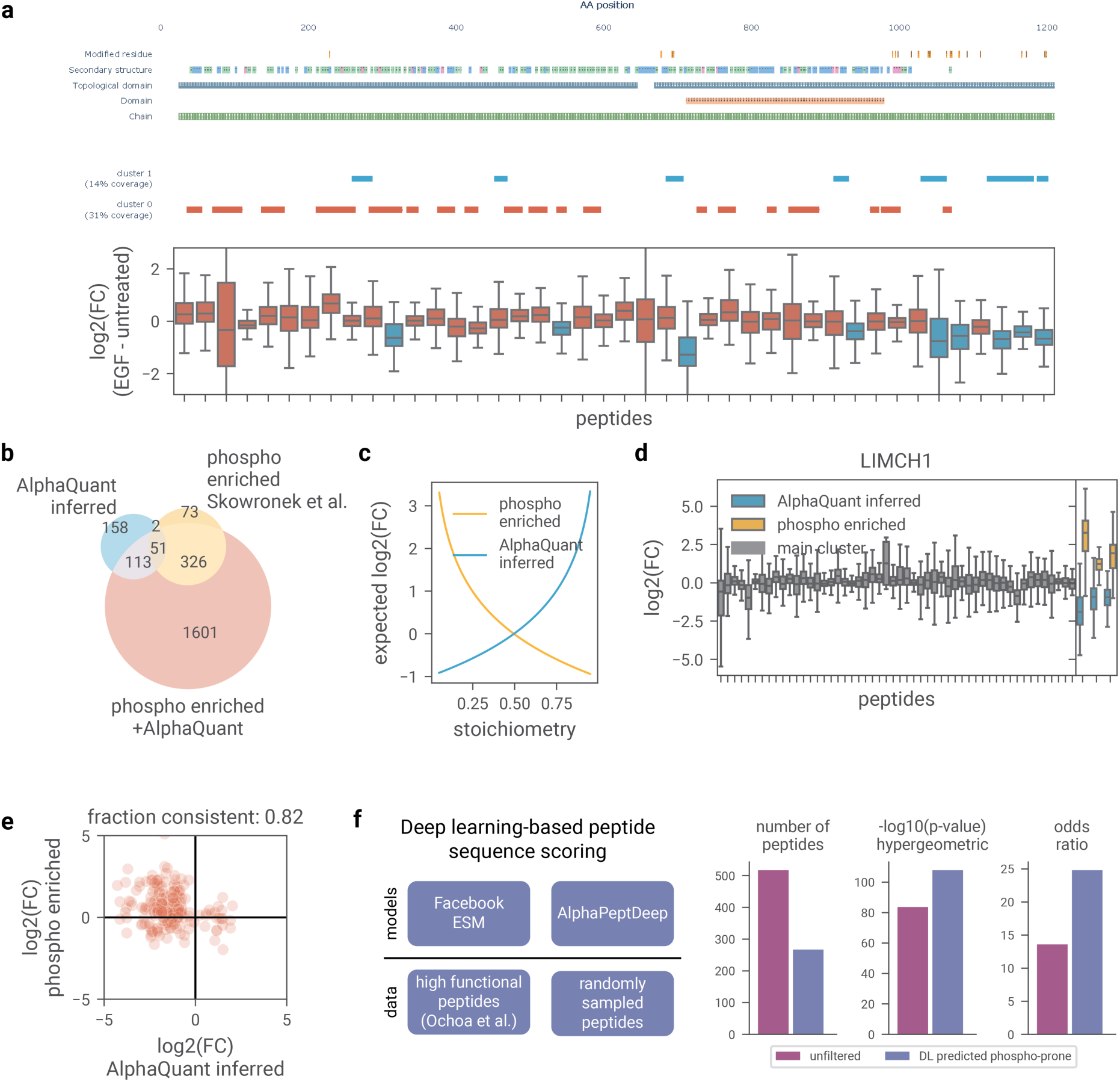
Analysis and validation of phosphopeptide detection. a) Outlier peptides detected by AlphaQuant mapped to the EGF receptor sequence. In the boxplot, peptides are ordered according to their position in the sequence and color-coded to match their corresponding clusters in the sequence plot. Adjacent peptides without sequence gaps appear as continuous strips in the sequence plot. b) Comparison of genes associated with regulated phosphopeptides: those identified through AlphaQuant analysis of unenriched data, those reported in the original phospho-enriched study, and those detected through AlphaQuant reanalysis of the enriched data. c) Simulation of stoichiometries of phosphopeptides vs. expected log2FC. d) Quantitative comparison of AlphaQuant-inferred and phospho-enriched peptides on LIMCH1. e) Correlation between fold-change directions in phospho-enriched data versus AlphaQuant-inferred data. f) A deep learning model for predicting functionally important phospho peptides from sequence features.

We validated our findings by comparing our results with a phospho-enriched dataset from previous EGFR stimulation experiments in our laboratory^37^. AlphaQuant analysis of this phospho-enriched dataset identified 2,091 genes with regulated phosphosites - a nearly five-fold increase from the 452 genes that we had reported before. As expected, most of the corresponding genes overlapped with those containing AlphaQuant-inferred outlier peptides (324 genes) (**Fig. 4b**). Although the overlap at the individual peptide sequence level was lower, it remained highly statistically significant (p < 10^−80^) (**Supplementary Fig. 1a**).

Note that phospho-enriched and AlphaQuant-inferred peptides are not expected to show complete overlap. Phospho-enrichment methods directly measure modified peptides while AlphaQuant quantifies their unmodified counterparts, leading to different sensitivities depending on modification stoichiometry. In phospho-enriched data, low stoichiometry modifications show larger log2FCs, whereas AlphaQuant-inferred outlier peptides exhibit stronger log2FCs for high stoichiometry modifications. These different properties can be simulated by comparing a fixed 0.5 stoichiometry reference state against variable stoichiometry conditions (**Fig. 4c**). Additionally, AlphaQuant-inferred peptides show a higher proportion of tyrosine (**Supplementary Fig. 1b**), a residue that presents enrichment challenges in general phosphoproteomics workflows.

For a direct dataset comparison, we first examined the protein Limch1 as an illustrative example (**Fig. 4d**). While most peptides remain unmodified, we identified three AlphaQuant-inferred outlier peptides and three peptides in the phospho-enriched dataset. Analysis of quantification between enriched and unenriched peptides revealed the expected counter-regulation pattern, in which increased phosphorylation corresponds to decreased unmodified peptide abundance. However, as shown in Fig. 5c, stoichiometric differences cause unmodified peptides not to decrease by the same amount that modified peptides increase. Across the entire dataset, approximately 80% of cases displayed the expected counter-regulation pattern (**Fig. 4e**). The remaining 20% might be due to noise in the measurement, or multiple different phosphosites for a given peptide, where the identified phosphosite is not the most abundant one. In accordance with this, additional filtering using our AQscore increased the fraction of consistent cases to 94% (**Supplementary Fig. 1c**).

**Figure 5:**
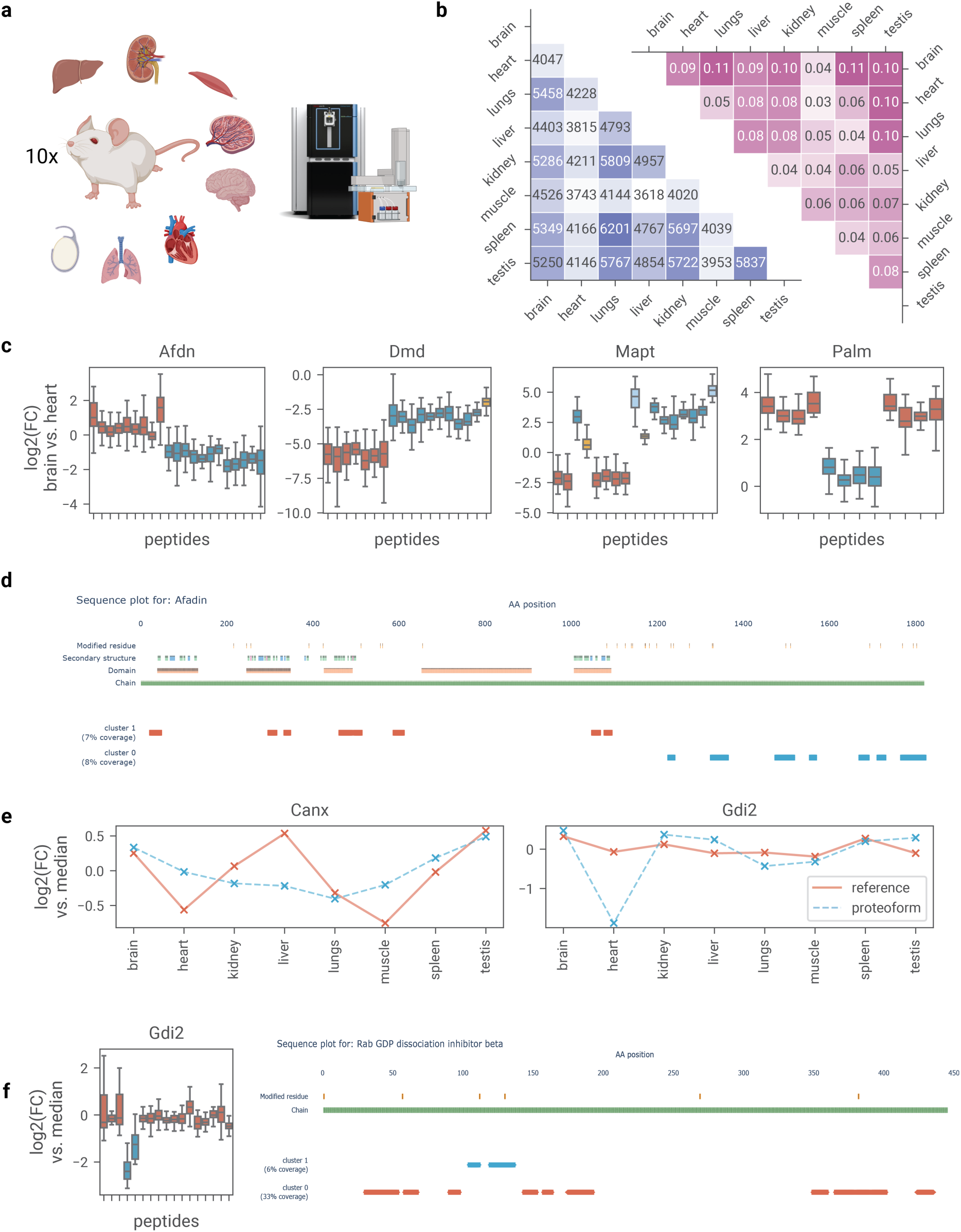
Proteoform analysis across mouse tissues. a) Tissues collected from 10 mice and measured on the Orbitrap Astral. b) Heatmaps quantifying detected proteoforms, showing proteoform frequency in each comparison (right) and genes accessible to testing (left). c) Representative examples of truncation and/or splicing in structurally important genes. Peptides are ordered along the sequence of the main isoform. d) Sequence alignment of Afdn. e) Tissue-specific proteoform profiles for Canx and Gdi2. f) Enhanced heart-specific phosphorylation of the Gdi2 gene.

In this phosphorylation-dominated dataset, we can generally attribute outliers to phosphorylation rather than any other modification, however, this may not always be the case. To have the possibility to still infer phosphorylation in datasets with multiple types of modifications, we developed a transformer-based binary classifier that combines our AlphaPeptDeep^38^ software with embeddings from the evolutionary scale protein language model (ESM, Meta/Facebook AI Research)^39^. That model was trained on a curated dataset of functionally significant phosphopeptides^40^ (**Fig. 4f, left**), enabling us to identify phospho-prone peptides in the human proteome (**Supplementary Fig. 2**). Restricting AlphaQuant outlier peptides to phospho-prone peptides retained more than half of the original outliers (5 times more than expected by chance). Notably, these filtered peptides were even more specific to the phospho-enriched dataset as evidenced by improved significance and odds ratio in hypergeometric testing (**Fig. 4f, right**).

### Proteoform centric analysis of a cross-tissue dataset

We evaluated AlphaQuant’s broader capabilities for detecting proteoforms using technical benchmarking with in silico proteoforms in the publicly available mixed species datasets^14^. This analysis demonstrated strong recall and precision within the expected range across thousands of proteins (**Supplementary Text**).

Encouraged by these results, we investigated tissue-specific proteoform variations. We measured eight mouse tissues (brain, heart, lung, liver, kidney, muscle, spleen, testis) using an Orbitrap Astral mass spectrometer with samples collected from ten mice (**Fig. 5a**), giving a comprehensive dataset with a total of 11,100 and an average of 7,700 quantified proteins and well separated tissues (**Supplementary Fig. 3**). After performing pairwise tissue comparisons in AlphaQuant, we counted proteoform occurrences using the AlphaQuant .proteoforms.tsv results tables. To ensure robust analysis, we required peptides to be detected in at least seven mice in both tissue pairs. The proportion of genes exhibiting proteoforms varied between 3-11% across tissue pairs (Fig. 5b), with brain and spleen showing the highest fraction of proteoforms and brain having the highest fraction of detected proteoforms.

For detailed investigation of AlphaQuant capabilities we compared brain and heart, two tissues frequently associated with alternative splicing. We focused on four genes exhibiting notable proteoform regulation in our dataset: Afadin (Afdn), Dystrophin (Dmd), Microtubule-associated protein tau (Mapt), and Paralemmin (Palm). Mapping peptides along their major isoform sequences revealed distinct clusters of sequence-adjacent peptides with consistent regulatory patterns (**Fig. 5c**). Detailed sequence analysis demonstrated clear separation between peptide groups based on protein regions, suggesting truncated or alternatively spliced proteoforms (**Supplementary Fig. 4a-c**).

Afdn and Dmd analysis indicated short proteoforms, with Afdn showing a particularly striking pattern - cluster 0 peptides aligned precisely with annotated protein domains, while cluster 1 displayed continuous coverage in the C-terminal region (**Fig. 5d**). Notably, none of the ENSEMBL^41^ GENCODE splice isoforms matched the observed pattern, suggesting either incomplete annotation or a different regulatory mechanism, such as proteolytic cleavage or degradation. Mapt exhibited complex splicing patterns indicative of multiple isoforms, consistent with reported Mapt splicing diversity^42^. In Palm, sequence alignment identified a distinct exon region with extensive peptide coverage spanning both exon junctions (**Supplementary Fig. 4d**).

AlphaQuant’s "median reference" analysis mode enabled us to compare individual tissues against a balanced median reference, facilitating global comparison across conditions and tracking of proteoform behavior across tissues. From the genes showing distinct proteoform profiles, we selected Canx and Gdi2 for detailed analysis. Both indicated regulation primarily through post-translational modifications rather than sequence-level changes like splicing or cleavage (**Fig. 5e**).

The endoplasmic reticulum chaperone protein Canx, which ensures proper protein folding and quality control, demonstrated clear tissue-specific regulation of the main protein. Analysis of its proteoform-associated peptides showed less pronounced abundance changes between tissues compared to the main protein. Mapping the peptides to the protein sequence identified a likely phosphorylation at the C-terminal end, including a MAPK3-targeted phosphoserine (**Supplementary Fig. 5a**). Differences were most pronounced in liver tissue, with an increase in the abundance of main protein peptides and a slight decrease in the abundance of peptides mapping to phosphorylation sites, indicating increased phosphorylation (**Supplementary Fig. 5b**). From this pattern, we infer that additional protein copies are directly phosphorylated, demonstrating tight co-regulation between protein expression and phosphorylation.

Gdi2, which regulates membrane trafficking through Rab protein control, maintained a stable main protein profile across tissues. However, two proteoform-associated peptides showed significant abundance decreases in heart tissue. AlphaMap sequence analysis revealed downregulation of two neighboring peptides that map to annotated phosphorylation residues, indicating enhanced phosphorylation of Gdi2 specifically in heart tissue (**Fig. 5f**).

Together, these diverse regulatory patterns spanning alternative splicing events and post-translational modifications demonstrate AlphaQuant’s capability to uncover tissue-specific proteoform regulation caused by multiple mechanisms.

## Discussion

Through tree-based quantification, AlphaQuant introduces innovations across four key elements of quantitative proteomics: differential quantification, missing values, quantitative accuracy, and proteoform inference. Implementation of differential quantification at the base ion level and its propagation along the tree demonstrates substantially increased recovery of regulated proteins. Integration of counting statistics with the tree-based model enables extensive and accurate recovery of proteins that are missing not at random, providing a statistically robust alternative to conventional imputation methods. Application of machine learning to tree-derived features, combined with acquisition metrics - peak shape quality, retention time, and identification probability - generates a quantitative accuracy score, addressing a major challenge in proteomics quantification. Using the principle of comparability within the tree, which relies on our dedicated statistical noise model, enables us to identify groups of co-regulated peptides, effectively addressing the protein grouping problem and enabling proteoform inference.

Beyond its core functionality, tree-based quantification provides flexible compatibility with diverse types of proteomics quantification and is applicable to both label-free and label-based methods. To advance proteoform analysis from bottom-up measurements, we address a key barrier in the field: the lack of standardized statistical pipelines for peptide-resolved proteoform analyses. Our software package seamlessly integrates proteoform investigation into standard differential quantification, offering this capability as an inherent feature of the workflow. We implemented AlphaQuant as a modular open-source Python framework, complete with a graphical user interface (GUI) and one-click installers, distributed under the permissive Apache License as part of the AlphaX ecosystem. It offers native support for standard output tables from multiple search engines, including AlphaPept^32^ and AlphaDIA^43^, as well as commonly used tools like MaxQuant^44,45^, FragPipe^46^, Spectronaut^47^, and DIA-NN^48^.

Bottom-up proteomics presents unique challenges for bioinformatic proteoform analysis. While algorithms can identify potential PTMs, they typically depend on enrichment methods to achieve high coverage, yet the number of actually occurring PTMs may still vastly exceed those detected. Additionally, these methods usually cannot resolve specific isoforms among multiple possible candidates. Quantification offers an elegant complementary approach by focusing on abundances of the unmodified peptides. This approach has inherently better proteome coverage and is agnostic to the type of PTM, however its method of detection is indirect and quantification biases might lead to false positives. While filtering reduced false positives in our benchmarking dataset, this led to decreased sensitivity. Additionally, the noise inherent in the data at peptide level, prevents the resolution of low stoichiometry PTMs. These issues may be addressed in the future by constantly improving peptide-level quantification accuracy and reliability.

As modern mass spectrometers achieve increasingly comprehensive peptide coverage^49^, the quantification-based approach holds promise for estimating global proteoform diversity in biological organisms. Analysis of our cross-tissue dataset detected approximately 1,600 genes with distinguishable proteoforms, most exhibiting three or fewer variants. Given the nature of the underlying dataset (bulk tissue analysis across different mice), many low-abundance isoforms likely remained undetected due to noise and limits in dynamic range. Nevertheless, these findings suggest that only a small subset of the vast set of theoretically possible proteoforms exists at detectable levels, consistent with previous observations in the field^50,51^.

Direct detection of modified peptides and splice peptides remains crucial, whether through enrichment methods or increasingly sensitive instruments and advanced bioinformatics approaches. Furthermore, peptide-resolved quantification tools can guide focused searches for modified peptides, both computationally and through targeted proteomics. We believe that proteoform-resolved statistical tools like AlphaQuant will help shift the paradigm from viewing proteins as simple point quantities toward embracing proteoform diversity and uncovering a new dimension of biological complexity.

## Methods

### Mixed species experiment

For the mixed species experiment for assessing quantitative accuracy, two different mixtures of HeLa tryptic digest (H, Pierce #1862824) and S. cerevisiae tryptic digest (Y, Promega V746A) with mixing ratios of 1:1, 1:10 and 1:100 were prepared (100 ng H : 100 ng Y, 100 ng H : 10 ng Y and 100 ng H : 1 ng Y).

### EGFR stimulation experiment

HeLa S3 cells (American Type Culture Collection) were cultured in high-glucose Dulbecco’s modified Eagle’s medium with 10% fetal bovine serum and 1% penicillin/streptomycin (all from Life Technologies, Inc). At a plate confluence of 80%, cells were treated for 10 min with 100 ng/ml animal-free recombinant human EGF (PeproTech) or Gibco distilled water (untreated, Thermo Fisher Scientific). Sample preparation was performed as previously described, similar to the in-stage tip protocol (Skowronek, 2022, MCP). In brief, the cells were washed three times with ice-cold TBS before lysis in 2% sodium deoxycholate in 100 mM Tris–HCl (pH 8.5) at 95 °C. Protein lysate was snap-frozen and stored at −80 °C. Protein reduction and alkylation was performed with 10 mM Tris(2-carboxyethyl)phosphine and 40 mM chloroacetamide and digestion with trypsin (Sigma–Aldrich) and LysC (WAKO) (1:100 dilution, enzyme/protein, w/w) at 37 °C overnight. Resulting peptides were purified by Sepax Extraction columns (Generik DBX) via styrenedivinylbenzene reverse-phase sulfonate (SDB-RPS), dried and reconstituted in a solution A∗ (0.1% TFA/2% acetonitrile (ACN)). Peptide concentrations were measured optically at 280 nm (Nanodrop 2000; Thermo Fisher Scientific). For extensive quantitative evaluation, different input amounts were injected (25, 50, 100, 200, 400ng).

### Mouse tissue collection

Six week old male mice of genetic background C57BL/6N were used for all analyses involving mouse tissue. The animals used were bred for scientific purposes, and the research in this project does not involve experiments on any animal, as defined by law. All animals were sacrificed prior to the removal of organs in accordance with the European Commission Recommendations for the euthanasia of experimental animals (Part 1 and Part 2). Breeding and housing as well as the euthanasia of the animals are fully compliant with all German (i.e., German Animal Welfare Act) and EU (i.e., Directive 2010/63/EU) applicable laws and regulations concerning care and use of laboratory animals.

After euthanasia with CO2, tissues were dissected and briefly rinsed in room temperature phosphate-buffered saline (PBS, pH 7.4). 1.5 mm punch biopsies of each organ were obtained and collected into a 96-well plate on dry ice.

### Sample preparation and protein extraction of mouse tissue samples

Tissue samples were collected in a BeatBox 96-well plate positioned on a magnetic stand to secure magnetic components. Subsequently, 100 µL lysis buffer (0.013% DDM in 60 mM TEAB) was applied to each well. The plate was sealed and homogenized using a BeatBox instrument at maximum output for 10 minutes. Following brief centrifugation, 80 µL of the homogenate was transferred to a fresh 96-well plate and the plate was sealed. Samples were boiled at 90°C for 60 minutes and subsequently allowed to cool to room temperature. After brief centrifugation, samples were sonicated using a Covaris instrument (300 seconds, 450 PIP, 50 DF, 200 CPB, 225 AIP), followed by centrifugation at 1000g for 2 minutes. Acetonitrile (ACN) was added to a final volume of 10% and the samples were then incubated at 75°C for another 60 minutes. The samples were allowed to cool to room temperature and centrifuged at 1000g for 2 minutes. Protein quantification was performed using a tryptophan assay. 10 μg dilutions of each sample were mixed and 0.1 µg of each trypsin and LysC was added. Digestion took place over night at 37°C. The reaction was quenched with trifluoroacetic acid (TFA) at a final concentration of 1% TFA. The samples were further desalted by 2x SDB-RPS (styrenedivinylbenzene reverse phase sulfonate) stage tips. For each sample, 20 µg of peptides were loaded onto the stage tips in triplicates and centrifuged at 800g until complete passage through the tips (approximately 10-15 minutes). The bound peptides were washed twice with 200 µL of wash buffer 1 (1% TFA in isopropanol) followed by a single wash with 200 µL of wash buffer 2 (0.2% TFA), with centrifugation at 1000g for approximately 10-15 minutes per wash step. Peptides were eluted into PCR collection tubes using 100 µL of elution buffer (1.25% NH₄⁺ in 80% acetonitrile) with initial centrifugation at 300g for 2 minutes, followed by additional centrifugation at 500g if needed for complete elution. The eluates were concentrated using a SpeedVac at 45°C for approximately 60 minutes. The dried peptides were reconstituted in 21 µL of EvoA buffer and resuspended at 2000 rpm at room temperature for 10 minutes. For samples processed with multiple stage tips, the peptides were pooled, and concentrations were determined using Nanodrop measurements with EvoA buffer as reference. A total of 200 ng of each sample was dispensed using an Opentrons OT- 2 automated liquid handling robot

### LC-MS/MS analysis of mixed species and EGFR stimulation experiment

Samples were measured using LC-MS instrumentation consisting of an EASY-nLC 1200 ultra-high-pressure system (Thermo Fisher Scientific), coupled to an Orbitrap Exploris 480 instrument (Thermo Fisher Scientific) using a nano-ES ion source (Thermo Fisher Scientific). Purified peptides were separated on a 50 cm in-house packed HPLC column with 75 µm inner diameter packed with 1.9 µm ReproSil-Pur C18-AQ silica beads (Dr. Maisch GmbH, Germany). Peptides were loaded in buffer A∗ (2% acetonitrile (v/v), 0.1% trifluoroacetic acid (v/v)) and eluted with a linear 95 min gradient of 3 and 5 to 30% of buffer B (0.1% formic acid, 80% (v/v) acetonitrile), respectively, followed by a 5 min increase to 60% of buffer B and 5 min increase to 95% of buffer B. The column was washed for 5 min at 95% buffer B and equilibrated for the next run at 5% buffer B for 10 min. The flow rate was kept constant at 300 nl/min. The column temperature was heated to 60 °C by an in-house developed oven containing a Peltier element. MS data were acquired in data-independent mode with MS1 automatic gain control target set to 300% in the 300 to 1400 m/z range with a maximum injection time of 45 ms and a resolution of 120,000 at m/z 200. Fragmentation of precursor ions was performed by higher-energy C-trap dissociation with a normalized collision energy of 30 eV. MS/MS scans were performed at a resolution of 15,000 at m/z 200 with an automatic gain control target of 1000%. The m/z range of 361 to 1033 was covered by 78 windows with 1 Da overlap (isolation windows of 9.6 m/z).

Technical replicates were acquired to evaluate reproducibility and quantitative accuracy by calculating CVs and mean of the replicate injections. Moreover, we alternated the MS run order to avoid potential carryover effects or any similar biases.

### LC-MS/MS analysis of mouse tissue samples

Digested peptides were loaded on EvoTip Pure C-18 tips (EvoSep) for subsequent LC-MS analysis. The samples were thawed at room temperature and centrifuged at 1000g for 2 minutes. Evotips were activated with a 3-minute incubation in 1-propanol. This was followed by a dual washing protocol using buffer B (0.1% formic acid in 99.9% ACN), with tips being centrifuged at 700g for 1 min after each wash. The activation process was repeated with another 3-minute 1-propanol treatment. The Evotips underwent two more sequential washes with buffer A (0.1% formic acid). For sample loading, each Evotip first received 70 μL buffer A prior to sample addition. The EvoTips were briefly centrifuged at 700g for 15 sec and the sample was subsequently added onto the present liquid. A washing step of the well containing the sample was performed by adding 10 μL buffer A to each well and transferring this wash to its corresponding tip. Peptide binding was facilitated by centrifugation at 700g for 2 min. Following a 50 μL buffer A wash, tips were covered with 150 μL buffer A and briefly centrifuged at 700g for 15 sec. Prior to LC-MS analysis, loaded tips were stored in an EvoTip box containing fresh buffer A.

Samples were measured on a Orbitrap Astral mass spectrometer (Thermo Fisher Scientific) coupled to an Evosep One liquid chromatography system (Evosep). An EASY-Spray source (Thermo Fisher Scientific) operating at 2200 V connected the two systems. Peptide separation was achieved using a Whisper Zoom 60 SPD gradient on an Aurora Rapid C18 UHPLC column (8 cm, 150 μm ID, 1.7 μm particle size, IonOpticks) at 50 °C. Sample acquisition was performed with field asymmetric ion mobility spectrometry using no FAIMS and exclusively in data-independent acquisition (DIA) mode. Orbitrap MS1 scans were recorded at a resolution of 240,000, a scan range from 380 to 980 m/z using 500% normalized AGC target and 3 ms maximum injection time. Isolation windows of 2 Th with a maximum injection time of 3 ms, 500% normalized AGC target, and 25% HCD collision energy were recorded.

### Processing of Public Mixed Species Benchmarking Datasets

The raw mass spectrometry data from the mixed species benchmarking datasets (**Fig. 2**) were obtained from PRIDE repository PXD005573. The large fold change (LFC) and small fold change (SFC) datasets were processed independently using Spectronaut version 14 (Biognosys) in library-free directDIA mode. Searches were performed against FASTA files provided by Spectronaut for *H. sapiens*, *S. cerevisiae*, *E. coli*, and *C. elegans*. All analyses were conducted using default software parameters.

### Processing of Public Phosphoproteomics Benchmarking Dataset

Pre-processed mass spectrometry data from the phosphoproteomics spike-in dataset were downloaded from PRIDE repository PXD014525 as a Spectronaut experiment file (.sne). Data tables were subsequently exported using the Spectronaut data export tool.

### Processing of Mixed Species Dataset for Accuracy Benchmark

Raw mass spectrometry data from the mixed species accuracy benchmark (**Fig. 3**) were analyzed using two search engines: Spectronaut^47^ version 18 (Biognosys) and DIA-NN^48^ version 1.8.2. For Spectronaut, analysis was performed using library-free directDIA mode, while DIA-NN analysis employed the "FASTA digest for library-free search" option. Both tools searched against UniProt reference proteomes (03/2021) for *S. cerevisiae* (UP000002311) and *H. sapiens* (UP000005640). Default parameters were used throughout, except for DIA-NN where high accuracy mode was enabled for QuantUMS^6^ quantification.

### Processing of EGFR Stimulation Dataset

The unenriched samples of the EGFR stimulation experiment were processed using Spectronaut version 17 in directDIA library-free mode against UniProt reference proteomes (03/2021) for *S. cerevisia* (UP000002311) and *H. sapiens* (UP000005640) using default parameters. The phospho-enriched samples were obtained from PRIDE repository PXD034128 and analyzed using Spectronaut version 18 with default settings, using a spectral library generated from phospho-enriched data available in the same repository.

### Processing of Public Mixed Species Benchmarking Dataset for Proteoform Regulation

Raw mass spectrometry data from the PRIDE repository PXD005573 was processed using DIA-NN version 1.9. Analysis was performed in high accuracy quantification mode, utilizing the "FASTA digest for library-free search" option with default settings otherwise. Reference proteomes were obtained from UniProt for four species: *S. cerevisiae* (UP000002311, 03/2021), *H. sapiens* (UP000005640, 03/2021), *E. coli* (UP000000625, 03/2021), and *C. elegans* (UP000001940, 07/2024).

### Processing of Mouse Dataset

Raw mass spectrometry data were analyzed using Spectronaut version 18 in directDIA library-free mode. Searches were performed against the *M. musculus* reference proteome from UniProt (UP000000589, 10/2023) using default software parameters.

### Base Ion Differential Quantification

Proteomics data is organized into a wide-format matrix where each row represents a base ion and each column represents an individual experimental run. For DIA data, these base ions can be fragments, precursors or peptide sequences. The data matrix is segregated by experimental conditions, and all intensity values are log2-transformed, with zero intensities encoded as NaN. To characterize technical and biological variation within conditions, log2FCs are calculated between samples of the same condition. These intra-condition comparisons are used to construct differential empirical error distributions (DEEDs) as previously described^9,52^. To account for intensity-dependent variance, DEEDs are generated separately for discrete intensity ranges.

Statistical significance of between-condition differences is assessed by comparing observed base ion log2FCs to their corresponding intensity-specific DEED, yielding empirical p-values. This approach is distribution-agnostic, as it directly models the underlying error structure from the experimental data rather than assuming a theoretical distribution.

For each base ion, N1 * N2 comparisons are performed, where N1 and N2 represent the number of samples in conditions 1 and 2, respectively. The resulting empirical p-values are converted to Z-values while preserving log2FC directionality. These Z-values are then aggregated and normalized using a scaled standard deviation derived from the corresponding covariance matrix^52^.

### Comparison of differential base ions

The relative regulation patterns between pairs of base ions are evaluated using the double differential empirical error distribution (DDEED) approach^52^. This method extends the single-condition DEED analysis by modeling the expected variation in log2FC differences between ion pairs. The DDEED is constructed by subtracting the respective DEEDs of two ions, thereby creating a noise model for log2FC differences.

To assess regulatory similarity between two base ions, log2FCs of log2FCs (FCFCs) are calculated by subtracting all log2FCs of the first base ion from those of the second base ion. These FCFCs are then compared against the appropriate DDEED to generate empirical p-values. The resulting p-values are transformed into Z-values and are summarized using a scaled standard deviation derived from the covariance matrix. The final p-value represents the probability of observing the measured differences under the null hypothesis of similar regulation between the two base ions.

### Between-sample normalization

Normalization between individual experimental samples is performed using a hierarchical single-linkage clustering approach described previously^5,9^, with samples being progressively merged based on their similarity. For each pair of samples, a log2FC distribution is constructed by calculating the log2FC for each base ion that is detected in both samples. Similarity is quantified by computing the variance of this log2FC distribution. When merging clusters, one cluster is rescaled by applying the median or mode of the log2FC distribution as a scaling factor. The merging process begins with individual samples as distinct clusters and iteratively combines the most similar clusters until all samples are unified in a single cluster.

For normalization within experimental conditions, samples were scaled using the median of the log2FC distribution as the normalization factor, as this provides a robust estimate of systematic shifts between samples. For between-condition normalization, a dual-metric approach is implemented: either the median or mode of the log2FC distribution is used as the scaling factor. The choice between median and mode is determined by their concordance - when the median and mode are similar, the median is selected; when they diverge significantly, the mode is chosen. This approach builds on the reasoning that the mode provides a more reliable estimate of the stable, non-shifting portion of the distribution in cases where substantial regulation occurs between conditions, though it is more susceptible to noise in its estimation.

### Constructing the tree

The hierarchical tree structure for each base ion is derived from the original search engine input tables. Each ion name encodes the complete path from the base ion to the root node. For example, in the identifier "SEQ_DALLVGVPAGSNPFR_MOD CHARGE_2_FRGION_b3_noloss_1", each component (i.e. SEQ, MOD, CHARGE, FRGION) represents a distinct hierarchical level.

The tree is assembled by creating nodes at each defined level of the hierarchy. Leaf nodes, representing the most granular level of the tree, are linked to their corresponding differential base ions. This hierarchical organization enables the systematic aggregation and propagation of statistical measures through the tree structure.

### Clustering along the tree

The clustering process begins at the parent level of base nodes and iteratively progresses upward through the tree. For each set of child nodes, pairwise comparisons are performed using their associated differential base ions, generating a comprehensive p-value matrix. These p-values are adjusted for multiple testing using the Benjamini-Yekutieli^53^ method, which was selected to account for the dependent nature of pairwise comparisons.

Clusters of similarly regulated nodes are identified using Ward’s hierarchical clustering algorithm. The significance threshold for cluster formation is set to 0.01 across most tree levels, with a more stringent threshold of 0.001 applied at the peptide level where proteoform inference occurs. At the peptide level, precursor quantities are used for clustering if available and clusters exhibiting minimal log2FC differences (default < 0.5) are merged to prevent over-segmentation. The main cluster for differential analysis is selected based on membership size. Following the calculation of precursor-level AQscores, clusters are reordered according to their cumulative AQscore, providing a quantitative basis for cluster prioritization.

### Propagating differential results up the tree

Z-values from grouped base ions are propagated upward through the tree structure by summarization and rescaling. At each level, the combined Z-value is scaled by the appropriate normalization factor, which is the square root of the number of contributing elements^9^. Z-values are computed by aggregating and rescaling the Z-values from their constituent child nodes.

To prevent statistical inflation from nodes with numerous measurements, constraints are applied during the propagation process. For precursors with multiple fragments, only the four base ions closest to the median Z-value are used in the differential analysis. Similarly, for proteins with four or more peptides, the least significant half of the peptides contribute to the final score.

The Z-value at the tree root is transformed into a p-value to generate the protein-level differential quantification result. These protein-level p-values are adjusted for multiple testing across all proteins using the Benjamini-Hochberg^54^ correction.

### Accuracy score model

We implement a gradient boosting regression model using scikit-learn’s HistGradientBoostingRegressor to predict absolute log2FC differences between precursor and protein log2FCs. The model undergoes hyperparameter optimization via RandomizedSearchCV with 10 iterations, exploring learning rates, tree depths, and sample thresholds. Optional feature selection reduces the feature space by 50% using f_regression scoring. The final model is evaluated through 3-fold cross-validation, with performance assessed via mean squared error and correlation coefficients. We extract multiple feature categories from each node in the tree. Basic node parameters included clustering metrics (fraction and number of main clusters), quality control metrics (consistency fraction, coefficient of variation, minimum intensity, and replicate count), and structural features (number of leaf and child nodes). For ’mod_seq_charge’ nodes, we calculate MS1-MS2 consistency. Additional cluster statistics include secondary cluster size, total cluster count, and between-cluster log2FC differences. Additionally, we add precursor-level features from the search engine output table, using all columns with relevant numerical information and averaging each value over all samples.

### Intensity based counting statistics

The statistical analysis of missing values employs intensity-based stratification to account for intensity-dependent detection patterns. For each intensity range, the empirical probability of missing values is determined from replicate measurements within each experimental condition. The analysis proceeds through a two-step statistical testing framework and all statistical tests are performed at the precursor level (mod_seq_charge) or at higher levels in the hierarchical tree (closer to the root).

First, the precursor with more detected values is tested for random distribution of missing values within its intensity stratum using a binomial test (randomness threshold: p > 0.1). If this randomness criterion is met, a second binomial test is performed on the condition with fewer detections. This test uses the intensity-specific missing value probability from the condition with more detections as the expected probability, evaluating the null hypothesis that the observed pattern of missing values in the lower condition occurs by chance.

When the initial randomness test fails (p ≤ 0.1), indicating systematic deviations from intensity-dependent random missing patterns also in the higher condition, Fisher’s exact test is applied as a more conservative, intensity-independent alternative.

Log2FCs are computed using the 10th percentile of observed intensities from the condition with more detections. The log2FC direction is assigned as negative in the direction of missing values, under the assumption that missing values represent lower abundance.

The resulting p-values are converted to Z-values and propagated up the tree structure as described above.

### Median reference analysis

To facilitate comparisons across multiple conditions, we construct median reference conditions using a systematic sampling approach. For each reference condition, we collect one base ion intensity from each experimental condition and calculate their median, creating a balanced reference point. Due to the presence of multiple samples per condition, we can in principle construct several distinct median reference base ions. The construction process is iterative: after creating each median reference base ion, the contributing base ions are removed from the pool to ensure independence of reference points. We only retain median base ions that are detected in at least 90% of all conditions. This procedure is repeated until no further valid combinations can be constructed given the remaining base ions and missing values in the samples.

### PTM site mapping

Site localization probabilities and their corresponding precursors are extracted from the Spectronaut results table. Modified peptides with overlapping sites are grouped after alignment against the protein sequence. Site classification uses a dual-threshold approach: sites with probabilities below 0.2 are classified as "likely not modified," while sites above 0.6 are classified as "likely modified." Complete linkage clustering is then performed to group compatible sites, ensuring that likely modified sites are never grouped with likely unmodified sites. After clustering, site probabilities are aggregated, and the sites with the highest probabilities are designated as the most likely PTM locations. Sites where the maximum probability falls below 0.6 are assigned NaN values to indicate uncertain localization, though these sites are still retained in the final output table.

### PTM to proteome normalization

To account for changes in protein abundance that might affect PTM measurements, differential expression results between phosphoproteome and unenriched proteome datasets are mapped by subtracting the log2FCs of the proteome data from the phospho data. The mapping between PTM sites and proteins is performed using gene names and UniProt synonyms to ensure robust matching between datasets.

A conservative significance adjustment approach for the phosphoproteome data is implemented by exponentially dampening p-values when the corresponding protein shows significant regulation in the same direction. The dampening factor is calculated as the ratio of the normalized log2FC to the original phosphosite log2FC, applied exponentially to the logged p-values. This results in order-of-magnitude decreases in significance even for small shifts in log2FC. In cases where protein regulation strength is equal to or stronger than the phosphosite regulation, or where normalization reverses the regulation direction, the p-value is set to 1.0. While heuristic, this approach provides a conservative estimate of phosphosite regulation by systematically down-weighting PTM changes that might be driven primarily by protein abundance changes.

The pipeline generates comprehensive results tables containing the adjusted log2FCs and p-values, with multiple testing correction performed using the Benjamini-Hochberg method. Summary statistics include the number of PTMs successfully mapped to proteins and the total number of proteins identified in both phosphoproteome and proteome datasets.

### Inference of functional phosphopeptides

For inference of potential functional phosphopeptides from sequence data, we implemented a two-stage deep learning approach combining protein language modeling with a specialized transformer architecture. In the first stage, protein sequences were encoded using the ESM-2^39^ model (Facebook/Meta AI Research) to generate contextual amino acid embeddings with dimensionality d=480. For each peptide, we extracted the corresponding subsequence embeddings from its parent protein’s embedding space, preserving the positional or structural context of the peptide within the full protein sequence.

The extracted peptide embeddings served as input to a custom transformer model (AlphaPeptDeep^38^) designed to predict phosphopeptide functionality. The model architecture consists of a 4-layer transformer encoder with dropout rate of 0.4, followed by a sequence attention pooling layer and a final linear classification layer. The model was implemented using PyTorch and contains approximately 11M trainable parameters.

For model training, we utilized the phosphopeptide functionality database from Ochoa et al.^40^, considering peptides with functionality scores ≥0.5 as positive examples. Negative examples were generated by in silico tryptic digestion of the human proteome (UniProt UP000005640_9606), selecting STY-containing peptides absent from the Ochoa database. The dataset was balanced through random subsampling of the UniProt peptides and split into training (80%) and test (20%) sets. The model was trained until convergence using cross-entropy loss, achieving precision and recall values of 0.93 and 0.99 respectively on the held-out test set.

The trained classifier was applied to score all STY-containing peptides in UP000005640_9606, generating a "function score" for each sequence. Peptides with function scores >0.5 were classified as "phospho-prone". Validation against PhosphoSitePlus^55^ database demonstrated highly significant concordance between predicted phospho-prone peptides and experimentally validated phosphopeptides (**Supplementary Fig. 2**). The resulting phospho-prone peptide annotations were integrated into the cluster analysis, with outlier clusters containing at least one phospho-prone peptide being labeled as phospho-prone.

### Benchmarking Differential Expression Analysis on Mixed-Species Datasets

AlphaQuant’s performance was evaluated across three distinct experimental datasets: a small fold change dataset, a large fold change dataset, and a phospho-peptide spike-in dataset. For all analyses, proteins were classified as significantly regulated when their protein-level FDR was below 0.05. In mixed-species samples, proteins from non-human organisms were considered true positives (regulated), while human proteins served as true negatives (non-regulated). Empirical FDR was calculated as the proportion of human proteins among all proteins identified as significantly regulated.

For the small fold change dataset analysis, AlphaQuant was configured to use protein identifiers as the tree root (spectronaut_fragion_ms1_protein) and identical housekeeping proteins for normalization as reported in the reference study^14^, with default settings otherwise. Results were compared against the statistical test outcomes published in the supplementary materials of the reference study, and the comparison was limited to proteins and precursors present in both datasets.

For the large fold change dataset, AlphaQuant was run in spectronaut_fragion_ms1_protein mode using the recommended housekeeping proteins for normalization. Two separate analyses were performed: one with the default setting excluding missing values (minrep_both = 2) and another where missing values were allowed (minrep_either = 2).

For the phospho-peptide spike-in dataset, a two-step analysis approach was employed. First, the Spectronaut data was separated by organism and each subset was processed independently using the AlphaQuant PTM site mapping module (modification_type="[Phospho (STY)]", perform_ptm_mapping=True, organism="human"/organism="yeast"). The processed tables were then merged and the combined dataset was analyzed using the AlphaQuant with default parameters. In parallel, the PTM site mapping formatted table was analyzed using directLFQ. For the directLFQ analysis, a two-group comparison using Welch’s t-test was performed for each protein, where at least two valid measurements per condition were required. Log2FCs between conditions were calculated, and p-values were adjusted for multiple testing using the Benjamini-Hochberg procedure.

### Benchmarking precursor level accuracy

AlphaQuant was applied to the Spectronaut output tables using protein identifiers as the tree root (spectronaut_fragion_ms1_protein), with default settings maintained for all other parameters. For DIA-NN analysis, the tree structure was extended to include QuantUMS-derived precursor intensities (diann_precursor_fragion_ms1_protein), as these were found to provide valuable additional information.

The main analysis was conducted using the AlphaQuant precursor results tables (results.prec.tsv) from the two highest abundance S. cerevisiae conditions (10ng S. cerevisiae vs. 100ng S. cerevisiae). AQscores were extracted directly from the corresponding column in the precursor results table. Quality scores for the search engine derived presented in **Fig. 3** were obtained from the DIA-NN and Spectronaut results tables.

### EGFR receptor stimulation analysis

For the unenriched 400ng EGFR stimulation dataset, AlphaQuant was run with functional phosphopeptide inference enabled (perform_phospho_inference=True) and default settings otherwise. The .proteoforms.tsv output for downstream analyses. The phospho-enriched dataset was processed using the AlphaQuant PTM site mapping module (modification_type="[Phospho (STY)]", perform_ptm_mapping=True, organism="human") and default parameters.

Visualization of peptide log2FCs and sequence patterns was performed using AlphaQuant’s built-in modules. The AlphaMapVisualizer module, a thin wrapper around the AlphaMap^36^ package, was utilized for sequence visualization, while peptide log2FCs were plotted with the FoldChangeVisualizer module.

For comparative analyses between datasets, gene identifiers were extracted from the PTM site ids, and Venn diagrams were generated to display the overlap.

Stoichiometry values were simulated across a range from 0.05 to 0.95. Expected log2FCs were calculated using two distinct approaches: for phospho-enriched samples, the direct ratio of stoichiometries was utilized, while for AlphaQuant-inferred measurements, the ratio of unmodified proportions (1 - stoichiometry) was calculated.

For log2FC comparisons between the datasets, a quality filter of 0.9 AQscore was applied to both datasets.

The enrichment of AlphaQuant-inferred phosphopeptides in the experimentally validated phosphopeptide dataset was evaluated using hypergeometric tests. The population size was defined as all unique peptides identified in the dataset, with the number of successes being peptides having both experimental PTM measurements and non-modified counterparts. Two separate analyses were conducted: one considering all AlphaQuant-inferred phosphopeptides, and another restricted to phospho-prone peptides. For each analysis, the sample size was determined by the number of inferred phosphopeptides in the respective category, with successes defined as the overlap between inferred and experimentally validated phosphopeptides. The enrichment strength was additionally quantified using odds ratios.

### Proteoform benchmarking analysis

The dataset was composed of four organisms with different log2FCs (*E. coli*, *C. elegans*, *H. sapiens* and *S. cerevisiae*), where *H. sapiens* stayed constant, *S. cerevisiae* was spiked in at a log2 ratio of -1, *E. coli* was spiked in at a log2 ratio of 1 and *C. elegans* at a log2 ratio of 2. Paired proteins were created by random assignment of *C. elegans* proteins to *E. coli* proteins. A shared identifier was generated for each *E. coli*-*C. elegans* pair, to which all peptides from both proteins were mapped, resulting in "artificial proteoforms."

The modified results table was processed with AlphaQuant using precursors as base ions (diann_precursor_protein), with a minimum requirement of two replicates per condition. True positives were defined as *E. coli*-*C. elegans* proteins, while all other proteins were designated as true negatives. Precision and recall were determined based on the classification of proteins with outlier clusters in the resulting .proteoforms.tsv table.

### Mouse tissue analysis

Two different analysis modes were employed with AlphaQuant on the mouse tissue dataset: a pairwise comparison between all tissues, and a comparison of all tissues against a median reference. In both comparisons, precursors were utilized as base ions (spectronaut_precursor_gene). For the pairwise comparison, a minimum threshold of 7 replicate values per condition was applied (minrep_both = 7), with default settings maintained otherwise. For the median reference comparison (multicond_median_analysis = true), different replicate thresholds were implemented: 7 replicates were required in the investigated condition and 3 replicates in the median reference (minrep_c1 = 7, minrep_c2 = 3), with default settings otherwise.

Proteoform fractions were calculated from the .proteoforms.tsv tables of the respective pairwise comparisons. The total number of genes in each table was determined, and the number of genes annotated with one or more outlier clusters was extracted. Outlier clusters coming from peptides that mapped to more than one gene were excluded. The fraction was then calculated as the ratio between these two values. Sequence patterns and peptide log2FCs were visualized using the AlphaMapVisualizer (based on AlphaMap) and FoldChangeVisualizer modules, respectively.

## Acknowledgements

We thank our colleagues in the department of Proteomics and Signal Transduction and the Clinical Proteomics group at the NNF Center for Protein Research for their help and fruitful discussions. We thank Patricia Skowronek, Marc Oeller, Juanjuan Wang, Georg Wallmann, and Julia Schessner for providing valuable feedback and datasets, Sophia Steigerwald and Tim Heyman for their assistance with MS data acquisition, Magnus Schwörer and Sander Willems for their contributions to code and computational infrastructure, Eugenia Voytik for work on visualizations and the graphical user interface, and Isabel Bludau for help with the integration of AlphaMap. This study was supported by The Max-Planck Society for the Advancement of Science and by the Bavarian State Ministry of Health and Care through the research project DigiMed Bayern (www.digimed-bayern.de).

## Author contributions

Conceptualization: C.A and M.M. Bioinformatic method development: C.A., WF.Z. Architecture of ecosystem algorithms & software: C.A., WF.Z., M.St. Experiments and proteomics data acquisition: M.T, C.A.M.W., E.H.R., F.R. Writing - original draft: C.A. and M.M. Writing - review and editing: all authors; Supervision: M.M.; Funding acquisition: M.M.

## Code availability

AlphaQuant is free software accessible under the permissive Apache license. It can be found at www.github.com/MannLabs/alphaquant

