## Supplementary Figures for "Tree-based quantification infers proteoform regulation in bottom-up proteomics data"

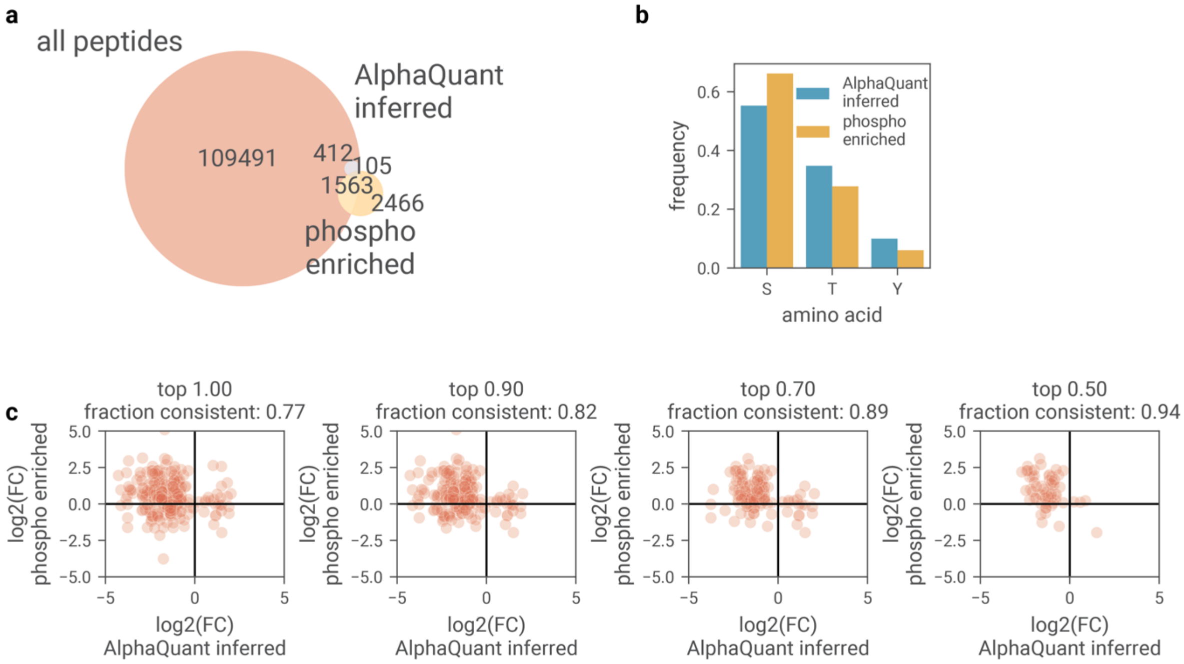


**Supplementary Figure 1: Comparison of AlphaQuant phosphopeptide inference against phospho-enrichment data.** a) Venn diagram showing the overlap between total peptides identified in unenriched samples, AlphaQuant-inferred regulated peptides, and phosphopeptides detected in phospho-enriched samples. b) Distribution of serine (S), threonine (T), and tyrosine (Y) phosphorylation sites among AlphaQuant-inferred peptides compared to experimentally validated phosphopeptides. c) Comparison of log fold changes (logFC) between EGF-stimulated and untreated conditions, comparing AlphaQuant-inferred phosphopeptides to experimentally validated phosphopeptides at increasing AQscore stringency thresholds (from no filtering to top 50% scores).

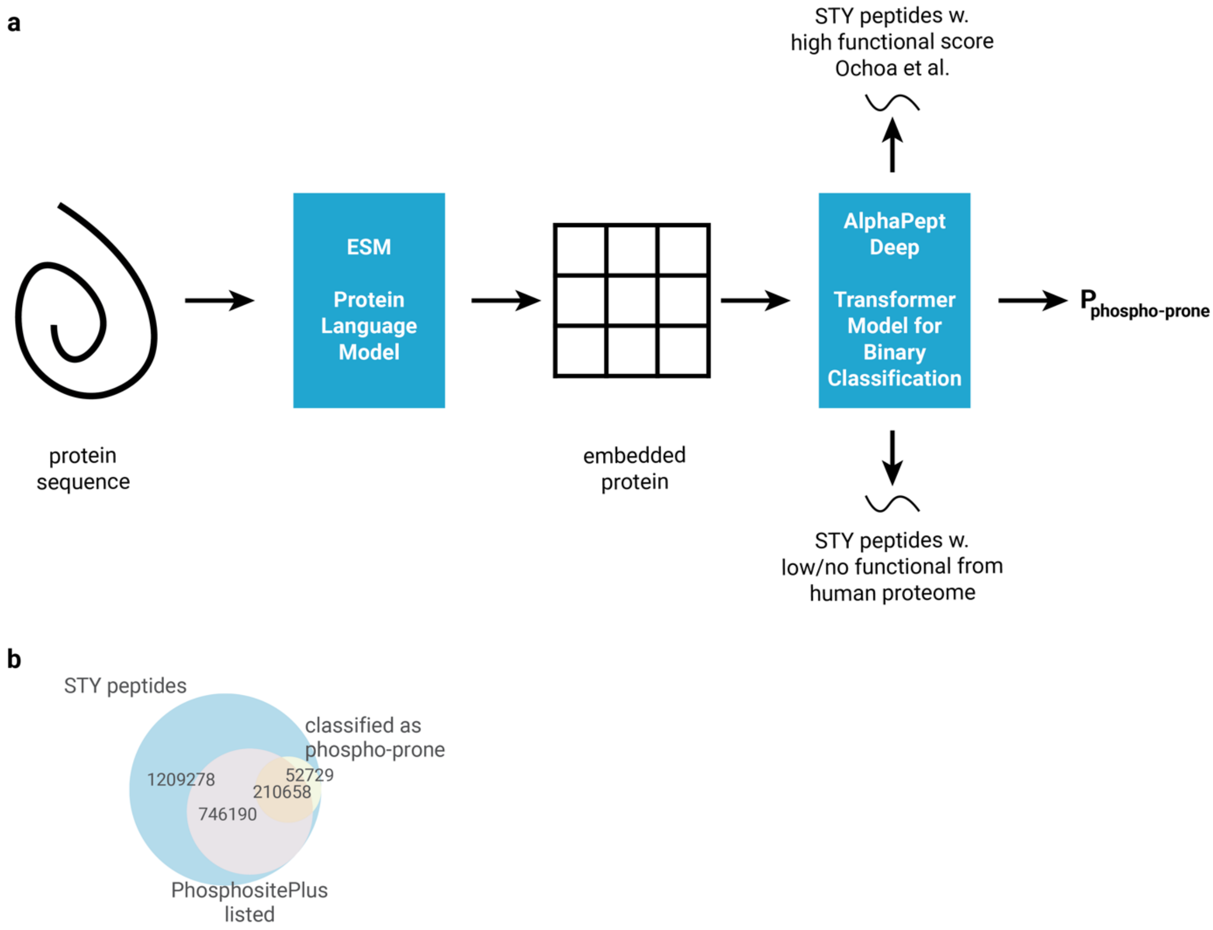


**Supplementary Figure 2: Concept and validation of the phosphopeptide classification model.** a) Model architecture: protein sequences are encoded using ESM-2[1], peptide embeddings are extracted maintaining positional context, and training uses phosphopeptides from Ochoa et al.[2] as positive examples and STY-containing tryptic peptides from UniProt[3] as negative examples. b) Overlap between STY-containing peptides from the human proteome, PhosphoSitePlus[4] database entries, and predicted phospho-prone sequences (P<10^-100, hypergeometric test).

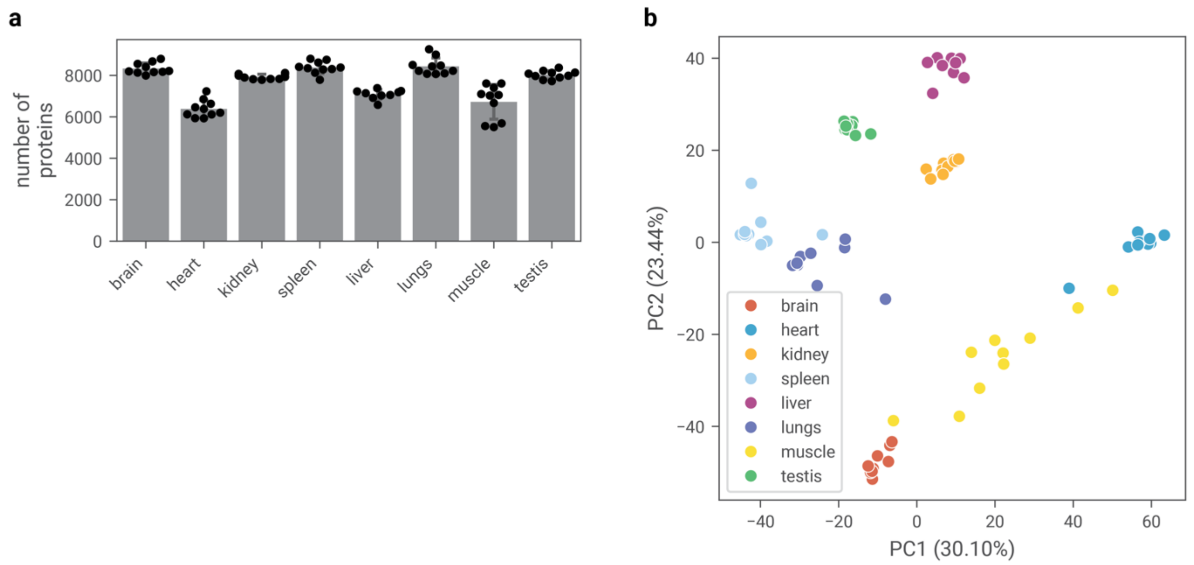


**Supplementary Figure 3: Overview of mouse tissue dataset.** a) Number of quantified proteins by directLFQ[5] per sample across different tissues. b) Principal component analysis of quantified proteins.

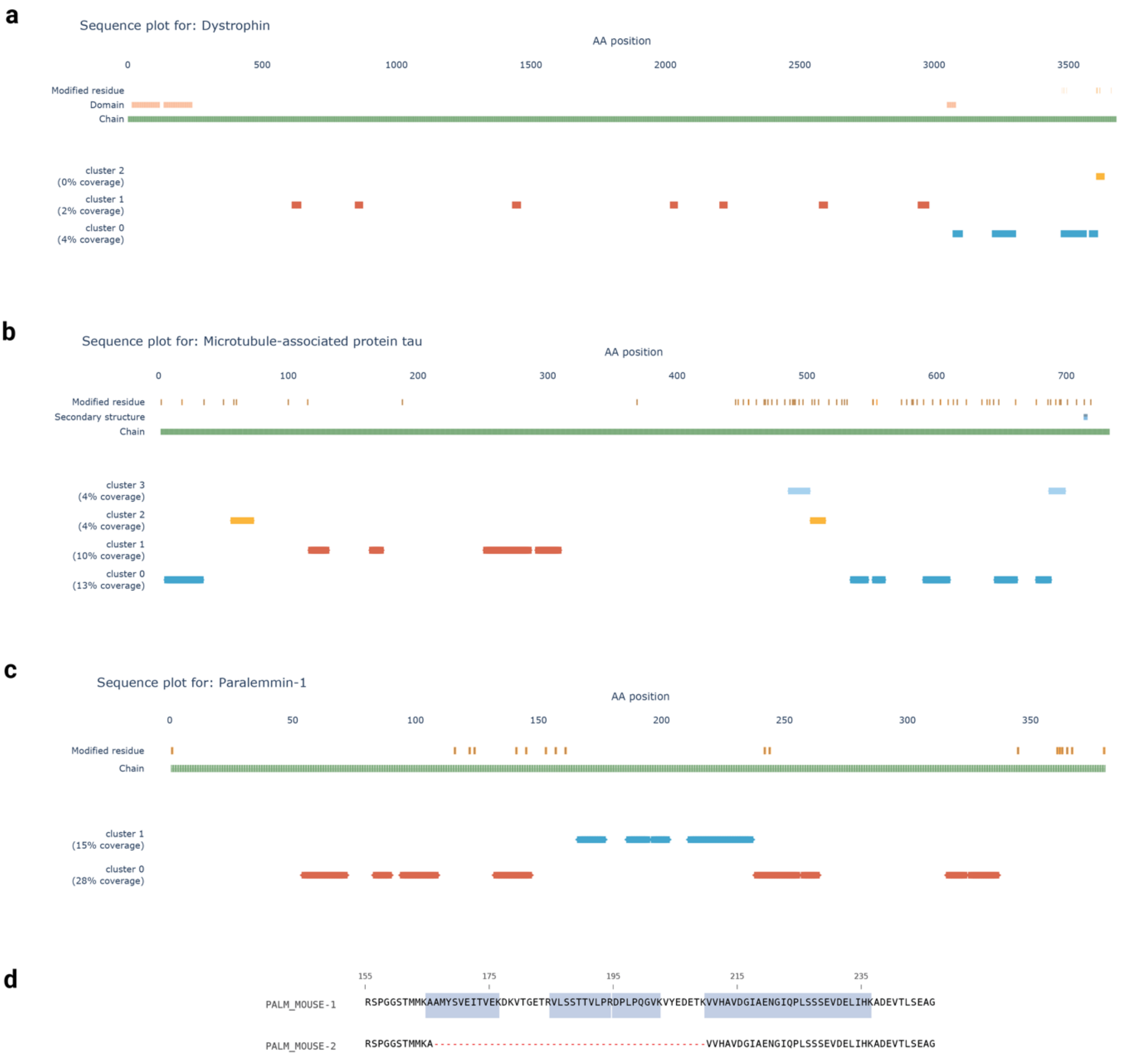


**Supplementary Figure 4: Peptide distribution analysis across main protein isoforms.** a)The Dystrophin protein (Dmd) exhibits two distinct peptide patterns in its main isoform: cluster 0 dispersed throughout the N-terminal region, followed by cluster 1 showing contiguous peptide alignment, and a single divergent peptide near the C-terminus. Cluster 1 comprises 10 peptides with adjacent alignment positions, appearing as continuous regions. b) The microtubule-associated protein tau (Mapt) exhibits four peptide clusters within its primary isoform. Clusters 0 and 1 are the most abundant, with cluster 1 containing two peptides that show no alignment with the canonical isoform sequence. The presence of these non-aligning peptides, combined with the distinct yet consistent peptide distribution pattern, indicates complex alternative splicing. c) In Parlemmin-1 (Palm), cluster 1 occupies a central position in the main isoform sequence, suggesting spliced out exons. d) Sequence analysis of the two relevant Palm isoforms (Q9Z0P4-1, Q9Z0P4-2) for this event, as documented in UniProt[3], reveals that cluster 1 peptides (marked blue) perfectly align with the annotated splice variant. None of the peptides marked is expected in Q9Z0P4-2, consistent with the different regulation observed.

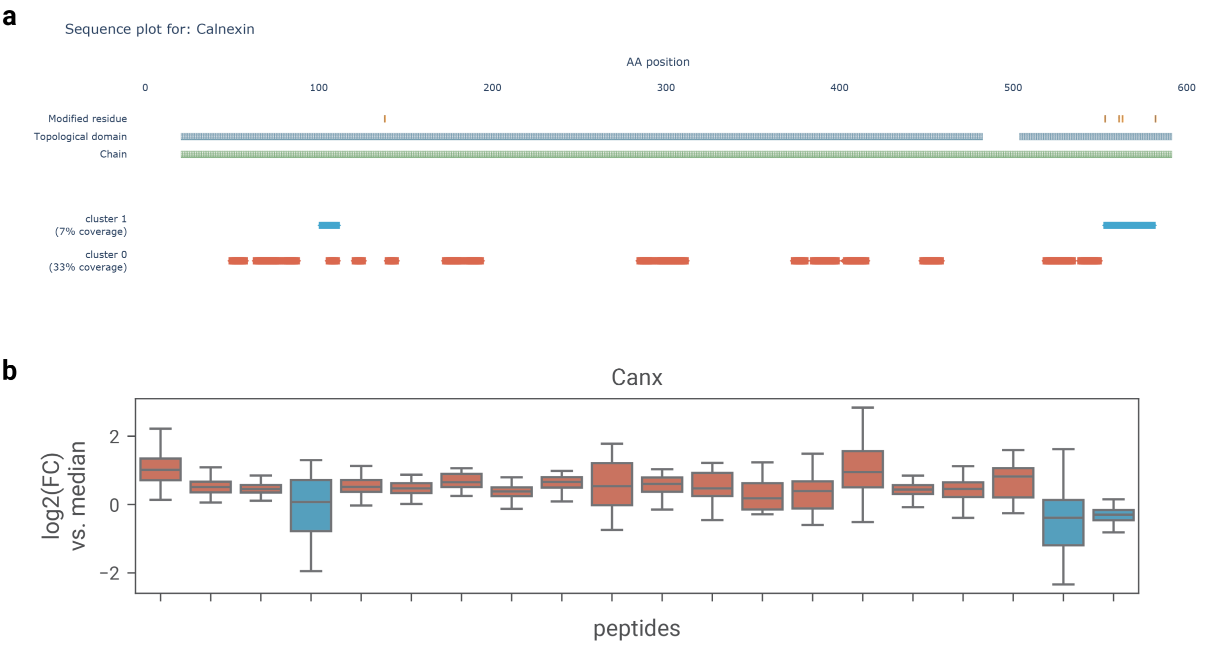


**Supplementary Figure 5: Sequence alignment and quantification of Calnexin (Canx) peptides.** a) Sequence alignment shows that C-terminal peptides of cluster 1 align with annotated modification sites, the second from the right being MAPK3-targeted phosphoserine. Peptides with adjacent alignment positions appear as continuous regions. b) Detail plots of peptides relative to the median reference show upregulation of cluster 0 and downregulation of cluster 1.
