## Supplementary Text for "Tree-based quantification infers proteoform regulation in bottom-up proteomics data"

### Benchmarking proteoform detection

AlphaQuant infers potential proteoforms by detecting clusters of similarly regulated peptides using its FCFC noise model. To evaluate proteoform detection accuracy, we designed a benchmarking dataset by modifying a mixed-species experiment containing proteins from four organisms spiked at distinct log2 ratios (*E. coli*: 1, *C. elegans*: 2, *S. cerevisiae*: -1, *H. sapiens*: 0). For each *E. coli* protein, we randomly assigned a *C. elegans* protein to create paired proteins. We then generated a unique protein identifier for each *E. coli*-*C. elegans* pair, ensuring all peptides from both proteins mapped to this shared identifier, thereby creating an "artificial proteoform." By converting only *E. coli* proteins to artificial proteoforms while preserving all other proteins in their native state, we established a benchmarking dataset that maintained the underlying data structure with minimal alteration (Fig 4a).

We analyzed this benchmarking dataset using AlphaQuant and examined its .proteoforms.tsv output to identify proteins classified as containing proteoforms (Fig 4b). The dataset comprised 700 "target" artificial proteoforms and 10,200 "decoy" native proteins. AlphaQuant successfully detected nearly all target proteoforms while reporting 180 (2%) of the decoys as potential proteoforms. Further inspection of these decoy hits revealed distinct regulation patterns in most cases, suggesting that systematic bias or false identifications contributed to this small fraction of false positives. By applying a quality score filter that summarizes the AQscore for each proteoform, we could eliminate the decoy hits. However, this filtering also removed many true positives, demonstrating a classical trade-off between specificity and sensitivity that users can adjust according to their analytical requirements.

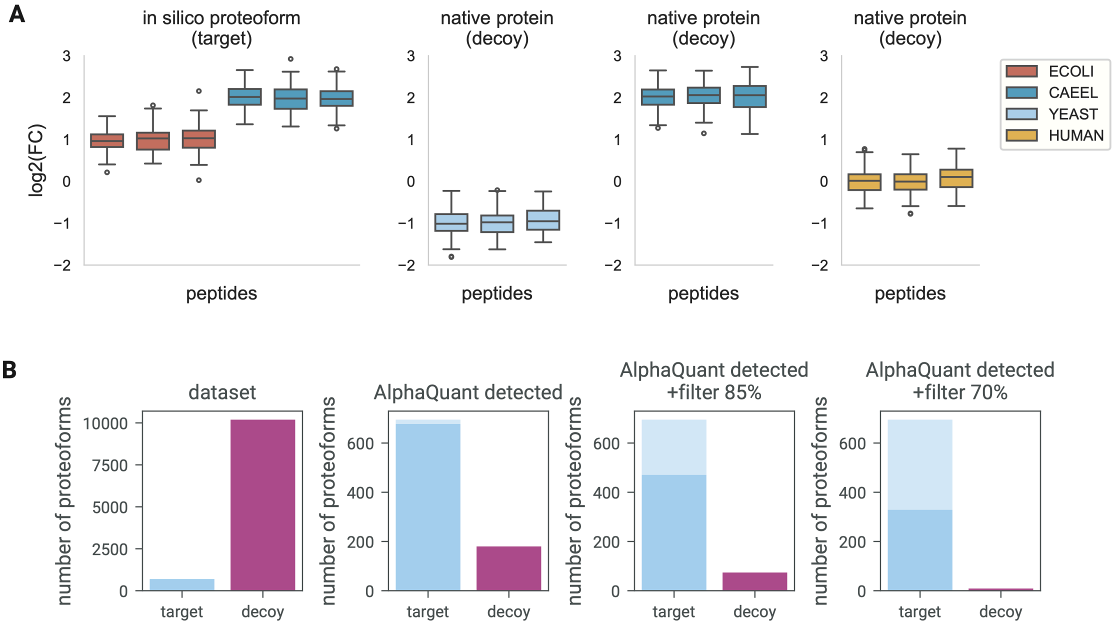


**Figure 1**: Constructing a benchmarking dataset for proteoform detection. A) Combining *E. coli* and *C. elegans* peptides in a mixed species dataset to create “target” proteoforms. Proteins from other organisms remain decoys. B) Distribution of targets and decoys in the dataset and recovery of targets and decoys with AlphaQuant.
